# Does a poor childhood associate with higher and steeper inflammation trajectories in the English Longitudinal Study of Ageing?

**DOI:** 10.1101/267559

**Authors:** Gindo Tampubolon, Maria Fajarini

**Affiliations:** Manchester Institute for Collaborative Research on Ageing, University of Manchester, Manchester, M13 9PL, UK; Evidence & Analytics, Manchester, M20 5WH, UK

**Keywords:** English Longitudinal Study of Ageing, childhood, C-reactive protein, fibrinogen, inflammation

## Abstract

Inflammation has been implicated in many diseases in later life of older Britons. Moreover, health outcomes in later life have also been markedly affected by childhood poverty. But no study has established whether childhood poverty has the effect of upregulating inflammation throughout later life. Using the English Longitudinal Study of Ageing (2004 – 2013) life history information and longitudinal observations of C-reactive protein and fibrinogen as inflammatory biomarkers, we studied the association between childhood condition and trajectories of inflammation for people aged 50 to 97 years. Retrospective childhood poverty some four to eight decades in the past was treated as a latent construct; attrition in longitudinal observations is addressed using inverse proportional to attrition weighting. The analytis revealed significantly higher levels of both biomarkers throughout later life among those with a poor childhood, though there is no evidence of a steeper inflammation trajectory among them. We discussed possible epigenetic changes underlying this strong and long arm of childhood condition. The results suggest that eliminating child poverty can prove to be a wise investment with the prospect of a lifelong reward.

## Introduction

Inflammation is implicated in many impaired functions and disease conditions in later life, including lung function, physical function, cognitive function, and type 2 diabetes^1,2^. Dysfunctions and diseases arising from such impairments make up a major proportion of the health care burden in an ageing population^3^. We have therefore studied inflammation in later life in Britain and examined how it is affected by the early stages of life course, particularly childhood condition.

Recently, the English Longitudinal Study of Ageing (ELSA) has been studied to gauge the contributions of later life inflammation and the life course on diabetes and it is found that together they associate with the onset of diabetes^2^. The life course score is constructed as the sum of indicators of parents’ socioeconomic position, individual education attainment, and individual socioeconomic position in midlife. A gradient of life course score is found with respect to the hazard of type 2 diabetes. Inflammation also contributes a significant effect on the hazard.

Another study of ELSA has shown that inflammation and coagulation contribute to cognitive deficits in older people who were observed repeatedly over an extended period in later life^1^. Both inflammation and cognitive systems are linked closely and have shared components^4^. Although some reports suggest that inflammation can be beneficia^5,6^, the study found that repeated high levels of C-reactive protein (CRP) and fibrinogen are associated with cognitive deficits. Another study of this longitudinal sample, focusing on multiple biomarkers of organ systems including inflammation, found that attrition in the sample is substantial. The study adjusted for attrition by using weighting since the attrition relates to the levels of inflammatory and other biomarkers^7^.

Given the accumulating evidence that inflammation is independently harmful to an array of health outcomes, what could be driving inflammation throughout later life? A recent work on ELSA suggests a temporally remote possibility: childhood poverty^8^. The study examined mental health, cognitive function, and gait speed in later life and found that a poor childhood, specifically lacking indoor toilet, fixed bath, running cold and hot water, and central heating, associates with more depression, poorer memory, and slower gait. In examining the association with childhood condition, which is potentially recalled inaccurately, the study used a latent construct of childhood poverty^8–10^. The study illustrates the necessity for care in handling information on childhood condition some four to eight decades in the past when explaining the effects of childhood condition on health outcomes in later life.

We aim to answer the question above by focusing on the intermediate step in the mechanism linking childhood condition through inflammation to dysfunction and disease: the link between childhood condition and repeated systemic inflammation. To focus our work we raise two questions: Does a poor childhood associate with repeatedly higher levels of CRP and fibrinogen? Does it also associate with a steeper increase in both biomarkers over time?

## Materials and methods

The English Longitudinal Study of Ageing (ELSA) is the main resource for a nationally-representative ageing study of the English older population. The first wave was in 2002 and subsequent waves follow biennially. Repeated biomarker information is available from the even numbered waves (2004/5, 2008/9 and 2012/3) when nurses visited the participants. The data are freely available from the UK Data Archive (www.data-archive.ac.uk.) as study number 5050. More details of the study are given elsewhere^7,11–13^.

We followed the literature^14^ which analysed ELSA’s sister study of Europeans in matching the sample with childhood condition to the sample with repeated inflammation information. This matching gives *N* = 3,821 at baseline, and *N* = 3,660 and 2,951 at subsequent waves. At baseline, the matched sample (compared to the unmatched sample) has relatively younger participants (65.7 *vs* 68.0 year), *t* = 10.2, *p* < 0.001, has lower CRP levels (2.5 *vs* 7.1 mg/l, *t* = 19.5,*p* < 0.001, and has lower fibrinogen levels (3.1 *vs* 3.4 g/l, *t* = 14.8, *p* < 0.001). Both samples have a higher proportion of women to men but there is no significant difference between the samples 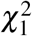 = 15.9, *p* < 0.26.

Following a study of ELSA and others^8,9^, the key exposure to childhood poverty is treated as a latent construct with potentially inaccurate indicators since these involve self-report of childhood condition. Self-report of childhood condition has been shown to be inaccurate in a British sample^15^, and a latent construct solution tailored to the ELSA sample has been applied^8^. Here a latent class analysis of childhood poverty, giving poor and non-poor latent classes, was applied to seven indicators including the sum of lacking (i) indoor toilet, (ii) hot and (iii) cold running water, (iv) central heating, (v) fixed bath;, plus (vi) overcrowding (more people than bedrooms), as well as (vii) number of books in the house, following our ealier work [16]. Notably, a latent class construct facilitates future cross-country comparison of childhood poverty based on recall information, since the indicators can be tailored to the situation in different countries.

Unlike the key exposure which is a latent construct, the key outcomes of inflammation (CRP and fibrinogen) were observed three times each. Blood samples were collected in three waves and kept deep-frozen until analysis at the Newcastle NHS hospital laboratory. Plasma samples were analyzed for fibrinogen using an ACL TOP CTS analyzer. The change in light transmission caused by the conversion of soluble fibrinogen in plasma to cross-linked insoluble fibrin is monitored by the analyzer, and the clotting time threshold is determined to be 37% of the total change. The clotting time is directly related to the concentration of fibrinogen in the plasma and this time is converted to concentration in g/l. Plasma samples were also analyzed for high sensitivity CRP, applying a particle-enhanced immunoturbidimetric assay, using Roche Modular P analyzer. CRP in the sample reacts specifically with anti-human CRP antibodies coated on the latex particles to yield insoluble aggregates. The turbidimetric absorbance of these aggregates is proportional to the CRP concentration in mg/l in the sample. Following the literature^16,17^, before the analysis we removed observations with CRP concentrations above 10 mg/l, indicating acute inflammation. Because CRP distribution is skewed, its log transform is used. Both biomarkers were used previously^1^.

We used mixed model to derive trajectories of both inflammatory markers; the model is variously known as latent growth curve model or random coefficients model. In applying mixed or random coefficients model, we tried both random intercepts and random slopes of age; but there were no significant variations in the slopes, so we settled with random intercepts. When modelling, in addition to age, sex, and wealth (in tertiles), we included covariates indicating socioeconomic positions earlier in the life course: education (threefold: less than high school as reference, high school, and college), and occupational class (threefold: routine manual as reference, intermediate, and managerial/professional).

Because the repeated observations have shrunk due to attrition^18^, we follow the extensive literature in using inverse proportional to attrition weighting^19–23^. Particularly following^20^, the attrition model includes age, sex, smoking, cognition, education, hypertension, cardiovascular disease, diabetes, and retirement status; then stabilised weights were computed with base model including age, sex, and education.

Lastly, because childhood poverty status used in the models is a predicted latent construct, its association with inflammation trajectories (while controlling for other covariates) was estimated using the new three-step method which adjusts the standard errors^24–26^. We presented two models for each biomarker: without and with interaction between age slopes and childhood poverty. If the interaction term is significant, then the childhood poor follow steeper (or gentler) trajectories of inflammation.

## Results

Women made up the majority of the sample (5,770; 55%) while the mean level of CRP is 2.475 mg/l (standard deviation [SD] 2.289 mg/l), of fibrinogen 3.139 g/l (SD 0.568 g/l), and of age 67 year (SD 9 yr); see Table 1.

**Table 1.** Description of analytic sample with reference categories in asterisks. Source: ELSA 2004-2013.

| Variable | mean or *N* | Std. Dev. or % |
| --- | --- | --- |
| CRP (mean, SD), mg/L | 2.475 | 2.289 |
| Fibrinogen, g/l | 3.139 | 0.568 |
| Age, year | 67.0 | 9.0 |
| Male* | 4,694 | 44.86 |
| Female | 5,770 | 55.14 |
| Marital status |  |  |
| Single* | 535 | 5.11 |
| Married | 7,239 | 69.18 |
| Separated/divorced | 1,105 | 10.56 |
| Widowed | 1,585 | 15.15 |
| Social class |  |  |
| Managerial | 4,140 | 39.56 |
| Intermediate | 2,532 | 24.20 |
| Routine manual* | 3,792 | 36.24 |
| Wealth |  |  |
| Poorest* | 2,810 | 26.85 |
| Middle | 3,660 | 34.98 |
| Wealthiest | 3,994 | 38.17 |
| Education |  |  |
| Less than high school* | 3,114 | 29.76 |
| High school | 4,174 | 39.89 |
| Some college | 3,176 | 30.35 |
| Overcrowding |  |  |
| No | 8,537 | 81.58 |
| Yes | 1,927 | 18.42 |
| Lacking facilities |  |  |
| None | 747 | 7.14 |
| 1 | 5,386 | 51.47 |
| 2 | 1,113 | 10.64 |
| 3 | 800 | 7.65 |
| 4 | 1,692 | 16.17 |
| 5 | 726 | 6.94 |
| Number of books |  |  |
| None or very few | 3,019 | 28.85 |
| One shelf | 2,438 | 23.30 |
| One bookcase | 3,192 | 30.50 |
| Two bookcases | 993 | 9.49 |
| Three or more | 822 | 7.86 |

The latent class analysis of childhood poverty revealed that 23.6% of the participants had a poor childhood at age ten (Table 2.). The indicators of childhood condition were lacking these facilities when growing up: indoor toilet, fixed bath, running cold and hot water, and central heating; an over-crowded house and few books in the house. The loadings showed for instance that having more books is negatively loaded on being a poor child whereas lacking more facilities is positively loaded on being a poor child.

**Table 2.** Loading of childhood poverty indicators. Source: ELSA 2004-2013.

| Poor, 23.6% | Loading |
| --- | --- |
| Intercept | 1.173 |
| Overcrowded | 0.139 |
| Lacking no facility | 0.000 |
| Lacking 1 | 2.555 |
| Lacking 2 | 1.514 |
| Lacking 3 | 1.654 |
| Lacking 4 | 2.783 |
| Lacking 5 | 2.217 |
| Num. books: none | 0.000 |
| One shelf | -0.797 |
| One bookcase | -1.264 |
| Two bookcases | -3.257 |
| Three or more | -4.315 |

## Trajectories of inflammation

Childhood poverty status was then used to predict trajectories of both inflammatory markers (CRP and fibrinogen) for people aged 50 to 97. The trajectories of both biomarkers showed annual linear increase, without tapering off or non-linearity. The data also did not support random age slopes, only random intercepts, giving each individual a different starting level at mid-life beyond those accounted for by observed covariates (sex, age, wealth, and lifecourse indicators of education and socioeconomic position). The trajectories increase annually by 0.003 (95% confidence interval [CI] 0.0003-0.005) mg/l for log of CRP and 0.03 (95% CI: 0.0015 −0.0043) g/l for fibrinogen (Table 3; complete table in Supplement). Those with a poor childhood have significantly higher levels of log of CRP and fibrinogen by 0.067 (95% CI: 0.003-0.124) mg/l and 0.039 (95% CI:0.003-0.075) g/l. The interaction terms in both slope models are not significant, suggesting no steeper increase in the inflammation trajectories among those with a poor childhood.

**Table 3.** Level and slope models for log of CRP (top pane) and for fibrinogen (bottom pane), adjusted for sex, wealth, education, occupational class (Complete table in Supplement). Source: the English Longitudinal Study of Ageing 2004 - 2013.

| Log CRP | Level model |  | Slope model |  |
| --- | --- | --- | --- | --- |
| | $\beta$ | *p* | $\beta$ | *p* |
| Poor childhood | 0.0673 | 0.04 | 0.3290 | 0.09 |
| Age | 0.0027 | 0.03 | 0.0117 | 0.91 |
| Age × poor childhood | | | 0.0039 | 0.17 |
| Fibrinogen | Level model |  | Slope model |  |
| | $\beta$ | *p* | $\beta$ | *p* |
| Poor childhood | 0.0394 | 0.03 | 0.0552 | 0.64 |
| Age | 0.0029 | < 0.001 | 0.0040 | 0.01 |
| Age × poor childhood | | | -0.014 | 0.41 |

Most importantly, the trajectories of both inflammatory markers are distinguished by childhood poverty status throughout later life (Figure 1). Beyond the annual increase in inflammation which applies to all, those whose first decade in life was marked by poverty showed distinctly higher levels of inflammation up to their ninth decade. There is no convergence beyond midlife once the course is set by poverty at ten. This is the first evidence of how long the reach of childhood poverty is (up to the ninth decade of life) and how deep (down to the levels of circulating inflammatory markers).

**Figure 1.**
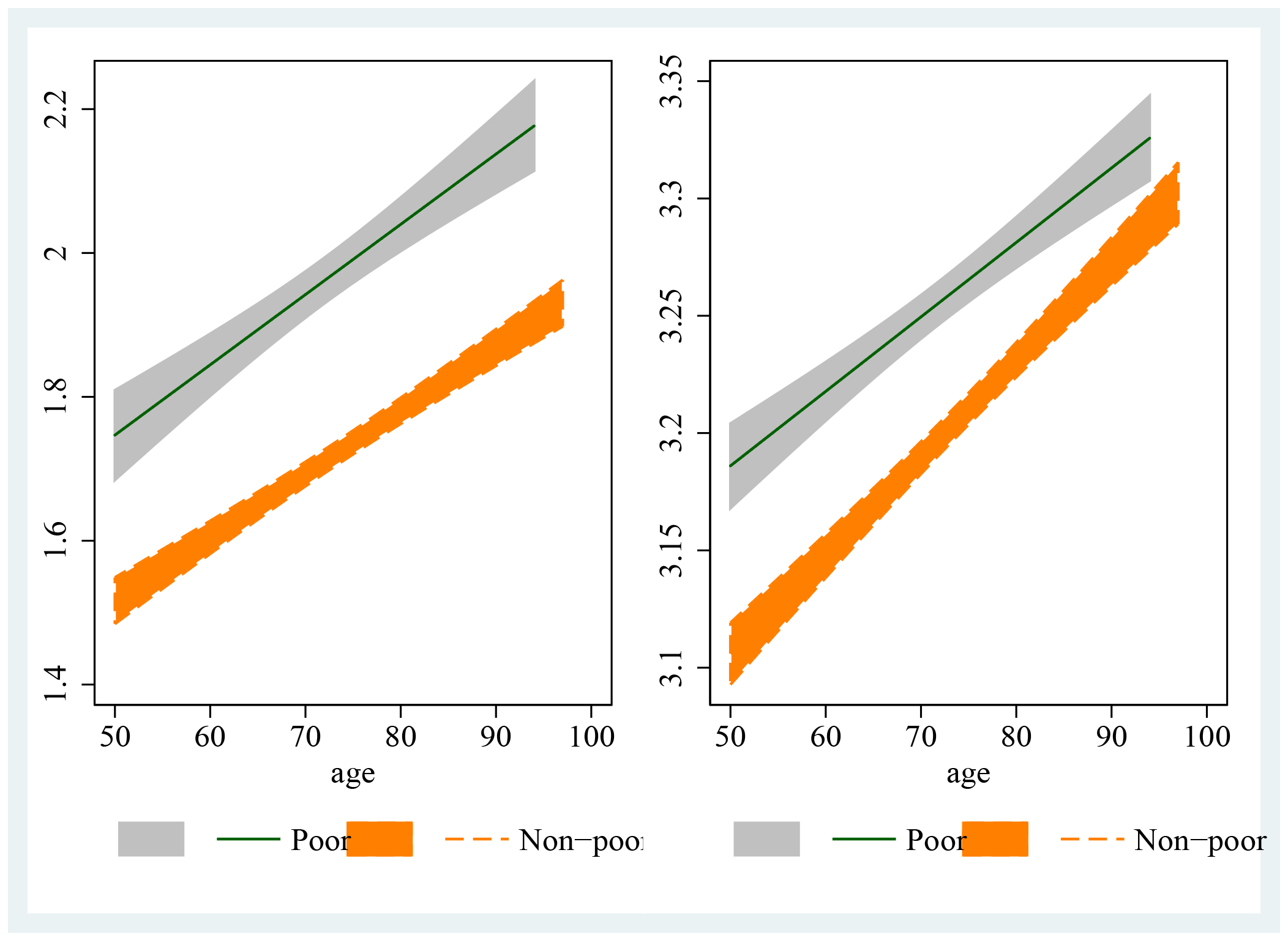
Trajectories of high sensitivity C-reactive protein (left pane) and fibrinogen (right pane), distinguished by childhood poverty (poor in grey, non-poor in orange) from 50 years onwards.

Both biomarkers showed stable and distinct poor versus non-poor trajectories throughout much of later life. Though there may be a discernible trend towards convergence between the gray and orange trajectories for fibrinogen (right pane), this is not statistically significant. Once poverty at ten puts some people on higher levels of fibrinogen at midlife, they track a different trajectory up to half a century later. On the other hand, we found no evidence that those with a poor childhood have a steeper inflammation trajectory.

## Discussion

We found that socioeconomic positions earlier in life, including education and occupation, predict levels of peripheral inflammation, as does contemporaneous wealth. Thus social inequalities in inflammation may be a driver to the widely observed social inequalities in health in later life^27^. But what drives these inflammation trajectories? An answer relates to a very early period in life. Being poor as a child not only goes with a higher level of inflammation at the onset of old age but throughout later life and without the prospect of convergence. Poverty in the first decade predicts inflammation until the ninth decade of life. The four near parallel mid-to-later-life inflammation trajectories, grouped by childhood poverty, showed the long arm of childhood condition and the depth of its reach ‘under-the-skin’ to affect inflammation function. Inflammation is not only key in later life, it is also set early in life.

Such a strong and lasting association can be explained with recourse to epigenetic changes, particularly DNA methylation, which imprints childhood material deprivation in the epigenome in a stable way. Material deprivation may compromise parental care, as parents struggle to provide for household members and lose valuable time for nurture. Such material deprivation may therefore impart social adversity to the child which can leave a biological mark^28–35^.

Because of the broader situation that material deprivation is a part of, it is not sufficent for an explanation to draw a mechanism from an animal model lacking the complexity of human social deprivation. However, non-human primate models, particularly macaques, can help. They have been used in a randomised design (of parental caring of frequent versus infrequent licking or grooming) to study the causal effect of early life deprivation on DNA methylation^36^. The study found increased DNA methylation patterns in the brain and the peripheral immune system. The changes are stable and organised, involving genes in the pathways of the immune system and the hypothalamic pitutitary adrenal (HPA) axis responsible for responding to stress. The peripheral immune system interacts with the HPA axis and has a role in brain function; evidence consistent with this interaction has been shown in this sample in our previous work^1^.

A key gene for regulating the HPA axis function, the glucocorticoid receptor (*NR*3*C*1), is activated in the hypothalamus in response to stress and releases glucocorticoid^28^. Glucocorticoid receptor is differentially expressed according to the experience of social deprivation, by epigenetic programming through histone acetylation and DNA methylation of the exon promoter^29,33^. This epigenetic programming differentiates similar DNA sequences phenotypically, resulting in blunted feedback by glucocorticoid and heightened stress response and demodulated immune system response, a pattern that is stable throughout the life course. The bidirectional interaction between the HPA axis and the immune system facilitates the imprinting of childhood adversity through epigenetic changes^33–35^.

This study has a number of weaknessess. First, by matching only individuals with childhood information with those with longitudinal observations, inevitablye some unmatched observations were set aside. It is impossible to measure the direction of possible bias this might entail. Second, although epigenetic changes are posited to be the mechanism, there is no direct evidence of the extent of DNA methylation in the collected blood sample. This is a potentially rectifiable weakness. Despite these weaknesses, this study has three major strengths. First, the sample is designed to represent the country and not only some clinical groups or regions, hence facilitating generalisation. Second, this is the first study to draw inflammation trajectories as they unfold with age. Finally, this study is also the first to link childhood condition (subject to recall error) with later life trajectories (subject to attrition), thus demonstrating the sustained link across the life course.

The effects of epigenetic changes have been postulated to be stable throughout the life course^33–35^. Here is supporting empirical evidence showing largely parallel inflammation trajectories between those with poor versus non-poor childhood. The effects of epigenetic change in childhood were not attenuated in later life. This is consistent with the view of a critical period in childhood, possibly through epigenetic imprinting^33–35^. In conclusion, the results suggest that eliminating childhood poverty can prove to be a wise investment with the prospect of a lifelong reward. It is never too early to invest in later life.

---

**Table 4. Supplement Table 1:** Log CRP: level model

|  | Coefficient | Standard Error | *p* value |
| --- | --- | --- | --- |
| Intercept | 0.5780 | 0.1083 | < 0.001 |
| Poor childhood | 0.0637 | 0.0310 | 0.040 |
| She | 0.0924 | 0.0260 | < 0.001 |
| Age | 0.0027 | 0.0012 | 0.026 |
| Middle wealth | -0.0755 | 0.0219 | < 0.001 |
| Wealthiest | -0.1370 | 0.0250 | < 0.001 |
| High school | -0.1223 | 0.0306 | < 0.001 |
| Some college | -0.2577 | 0.0374 | < 0.001 |
| Intermediate | -0.0747 | 0.0334 | 0.025 |
| Managerial | -0.0662 | 0.0325 | 0.042 |
| Married | 0.0109 | 0.0582 | 0.85 |
| Separated, divorced | -0.0477 | 0.0672 | 0.48 |
| Widowed | 0.0274 | 0.0635 | 0.67 |
| SD random int. | 0.7436 | 0.0095 | < 0.001 |

**Table 5. Supplement Table 2:** Log CRP: slope model

|  | Coefficient | Standard Error | *p* value |
| --- | --- | --- | --- |
| Intercept | 0.7884 | 0.1887 | < 0.001 |
| Poor childhood | 0.3290 | 0.1954 | 0.092 |
| She | 0.0921 | 0.0261 | < 0.001 |
| Age | -0.0003 | 0.0025 | 0.91 |
| Age × Poor childhood | 0.0039 | 0.0028 | 0.17 |
| Middle wealth | -0.0756 | 0.0219 | < 0.001 |
| Wealthiest | -0.1374 | 0.0250 | < 0.001 |
| High school | -0.1229 | 0.0306 | < 0.001 |
| Some college | -0.2583 | 0.0375 | < 0.001 |
| Intermediate | -0.0748 | 0.0335 | 0.026 |
| Managerial | -0.0647 | 0.0326 | 0.047 |
| Married | 0.0100 | 0.0588 | 0.86 |
| Separated, divorced | -0.0509 | 0.0680 | 0.45 |
| Widowed | 0.0276 | 0.0642 | 0.67 |
| SD random int. | 0.7433 | 0.0095 | < 0.001 |

**Table 6. Supplement Table 3:** Fibrinogen: level model

|  | Coefficient | Standard Error | *p* value |
| --- | --- | --- | --- |
| Intercept | 3.0722 | 0.0623 | < 0.001 |
| Poor childhood | 0.0394 | 0.0184 | 0.032 |
| She | 0.1059 | 0.0151 | < 0.001 |
| Age | 0.0029 | 0.0007 | < 0.001 |
| Middle wealth | -0.0387 | 0.0146 | 0.0081 |
| Wealthiest | -0.0595 | 0.0162 | < 0.001 |
| High school | -0.0301 | 0.0180 | 0.094 |
| Some college | -0.1078 | 0.0216 | < 0.001 |
| Intermediate | -0.0263 | 0.0196 | 0.18 |
| Managerial | -0.0184 | 0.0189 | 0.33 |
| Married | -0.0443 | 0.0312 | 0.16 |
| Separated, divorced | -0.0569 | 0.0371 | 0.13 |
| Widowed | -0.0208 | 0.0352 | 0.55 |
| SD random int. | 0.3898 | 0.0074 | < 0.001 |

**Table 7. Supplement Table 4:** Fibrinogen: slope model

|  | Coefficient | Standard Error | *p* value |
| --- | --- | --- | --- |
| Intercept | 2.9970 | 0.1129 | < 0.001 |
| Poor childhood | 0.0552 | 0.1181 | 0.64 |
| She | 0.1060 | 0.0151 | < 0.001 |
| Age | 0.0040 | 0.0015 | 0.0088 |
| Age × Poor childhood | -0.0014 | 0.0017 | 0.41 |
| Middle wealth | -0.0385 | 0.0146 | 0.0085 |
| Wealthiest | -0.0592 | 0.0162 | < 0.001 |
| High school | -0.0299 | 0.0180 | 0.096 |
| Some college | -0.1077 | 0.0216 | < 0.001 |
| Intermediate | -0.0260 | 0.0196 | 0.18 |
| Managerial | -0.0185 | 0.0188 | 0.33 |
| Married | -0.0438 | 0.0312 | 0.16 |
| Separated, divorced | -0.0558 | 0.0372 | 0.13 |
| Widowed | -0.0207 | 0.0352 | 0.56 |
| SD random int. | 0.3899 | 0.0074 | < 0.001 |

